# The Dilemma Of Disobedience: An ERP study

**DOI:** 10.1101/2021.07.05.451127

**Authors:** Eve F. Fabre, Mickaël Causse, Maryel Othon, Jean-Baptiste Van Der Henst

## Abstract

The present experiment aimed at investigating the decision-making and the associated event-related potentials (ERPs) of subordinates under hierarchical pressure. Participants (N = 33) acted as UAV operators and had to decide to crash their defective drone either on a civilian site killing all civilians present on the site or on a military site destroying military material but preventing any human losses. While in the no-command condition, participants decided according to their own preferences, in the command condition they were ordered to protect the military material at the expense of civilians for undisclosed strategic reasons. The results revealed that in the no-command condition participants almost always crashed the drone on the military site (96%), whereas in the command condition they chose to obey orders and sacrifice civilians to protect the military material 33% of the time. In the command condition, participants were longer to make their decisions, mobilizing greater attentional and cognitive resources (i.e., greater P300 responses) to resolve the conflict between their internal moral values and the orders they were given (i.e., greater N200 responses) than in the no-command condition, where they automatically applied the “you shall not kill” rule. Participants also showed a greater negative affective response (i.e., greater P260 amplitudes) after choosing to disobey than to obey orders. This result suggests that disobeying authority could be perceived as a greater moral violation than obeying and sacrificing civilians, suggesting that individuals may sometimes choose to obey malevolent authority to avoid the negative affective reaction triggered by disobedience.

## 1. INTRODUCTION

In the early hours of 26 September 1983, alarms from a secret command center near Moscow indicated that Soviet satellites had detected that various ballistic missiles had just been launched from an American base. Stanislav Petrov was a lieutenant colonel in the Soviet air defense forces and was the duty officer that night. Petrov, who doubted the reliability of this new warning system and knew that reporting the event (as he had to) would have triggered a full nuclear retaliation from USSR army, made the decision to violate the protocol and to wait for ground radar confirmation. Eventually, Petrov was proven right (the satellite had mistaken the sun’s reflection off the tops of high-altitude clouds for a missile launch) and his decision to disobey orders that night is thought to have prevented a nuclear world war and the death of hundreds of millions (if not billions) of people.

### 1.1. The Dilemma of (Dis)Obeying to Authority

The story of Stanislav Petrov is an extreme illustration of the struggle endured by subordinates when they have to decide whether to obey or not poor and/or immoral orders. In the ’60s, Stanley Milgram was the first to investigate the impact of malevolent authority on subordinates’ decision-making (Milgram, 1963). From a large set of studies, he concluded that obedience is ingrained in most of us and that ordinary people are very likely to follow orders given by an authority figure, despite the dramatic consequences this behavior may have (Milgram, 1963; 1965; 1974). Milgram (1974) proposed various explanations to the fact that most people obey malevolent authority. A first assumption was that coercion triggers a change in subordinates’ *sense of agency* (i.e., “the subjective experience of controlling one’s actions and, through them, events in the outside world”, Haggard & Tsakiris, 2009; Pfister et al., 2014), which may make them feel less responsible for their actions and decrease their empathy for the victim. The results of two recent neuroscience studies support this hypothesis (Caspar et al., 2016; 2020). A second assumption Milgram made in his paper entitled *The Dilemma of Obedience* (Milgram, 1974) was that authority may also affect subordinates’ moral sense:

The force exerted by the moral sense of the individual is less effective than social myth would have us believe. […] Even the forces mustered in a psychology experiment will go a long way toward removing the individual from moral controls.

When subordinates are instructed by their hierarchy to execute harmful orders, they are put in the situation of having to choose between two competing internal values: “*you shall obey orders*” and “*you shall cause no harm*” (Haidt & Graham, 2009). If they choose to disobey orders, they act in accordance with their moral values but risk being punished for disobeying. If they choose to obey, they ensure that they will not suffer retaliations from their superior, but have to act against their values. In order words, subordinates face a dilemma, in which they are given the choice between two undesirable alternatives (Lotto et al., 2014; Sarlo et al., 2012). Moral dilemmas have been widely used in both behavioral and neuroscience studies to investigate the way affective and cognitive processes shape moral decision-making (e.g., Christensen & Gomila, 2012; Greene at al., 2001; Greene, 2015). However, to our knowledge, the dilemma to obey or not malevolent orders has received little attention. The present study aimed at filling this gap using event-related potentials (ERPs) methodology and investigate whether being given orders affects the subordinates’ moral controls.

### 1.2. The ERP Studies on Moral Dilemmas

The analysis of ERPs allows to investigate the attentional, cognitive and affective processes underpinning decision-making with a high temporal resolution (Fabre et al., 2015; Ibanez et al., 2012; Luck & Kappenman, 2012) and is particularly adapted to the study of moral decision-making (Wagner et al., 2017). In the last decade, more than a dozen studies investigated the brain electrophysiological correlates of both moral judgement (Leuthold et al., 2015; Peng et al., 2017; Yang et al., 2013; Yoder & Decety, 2014) and moral decision-making (Chen et al., 2009; Pletti et al., 2015; Sarlo et al., 2012, 2014; Yun et al., 2019; Zhan et al., 2018; 2020). In moral decision-making studies, participants are presented with moral dilemmas depicting a situation where one is given the choice between two undesirable alternatives (Lotto et al., 2014; Sarlo et al., 2012), serving as an illustration of the tension between different normative ethical theories (e.g., Foot, 1967; Reynolds et al., 2019). In most of these studies (i.e., Pletti et al., 2015; Sarlo et al., 2012, 2014; Yun et al., 2019; Zhan et al., 2018; 2020), moral dilemmas were presented in written form in a series of slides, with the first slide presenting the scenario, the second slide and the third slide presenting respectively the first option “A” and the second option “B”, and finally, a decision slide where the letters A and B were displayed.

The results of these experiments revealed three main ERPs associated with the onset of the decision slide: the N200, the P260 component and the Late Positivity Potential (LPP) component. The N200, a negative deflection peaking in the 200 – 300 ms time window and maximal at medial and frontal sites (Gehring, & Fencsik, 2001; Kimura et al., 2013), was found in one study (Yun et al., 2019) and is functionally interpreted as reflecting the internal moral conflict during the decision-making. The P260 is a positive component belonging to the family of P2-components occurring in the 200 – 350 ms time window after the stimulus onset and maximal at frontal-central sites (e.g., Sarlo et al., 2012). The P260 is functionally interpreted as reflecting an aversive affective reaction and was found to be positively correlated to the to the unpleasantness experienced during moral decision-making (Sarlo et al., 2012; 2014; Pletti et al., 2015; Zhan et al., 2018; 2020). Finally, the LPP is a positive component that occurs in the 300 – 600 ms time window after the stimulus onset and maximal at central–parietal sites (Luck & Kappenman, 2011). This component was functionally interpreted as reflecting the allocation of attentional resources and the cognitive effort to resolve moral conflicts (Sarlo et al., 2012; 2014; Pletti et al., 2015; Zhan et al., 2018; 2020).

### 1.3. The present study

The present experiment aimed at investigating both the cognitive and the affective processes underpinning decision-making under hierarchical pressure and the associated ERPs in a military-like context. The military context is particularly adapted to the study of obedience to authority (Crosbie & Kleykamp, 2018; Wolfendale, 2009), both because military organizations are characterized by their strong hierarchical structure (Baarle et al., 2015) and because the duty of military obedience is very likely to conflict with one’s internal values (Gaeta, 1999; Frederick, 2010). As disobedience is extremely rare in military servants, we chose to conduct the experiment on civilian participants (Collart et al., 2015). The latter had to imagine that they had been recruited by the army as civilians to operate uninhabited aerial vehicles (UAV, also called drones) in a war zone, with the aim of collecting intelligence on enemy locations and guiding bombing runs – as it is already the case in the United States Air Force (Hennigan, 2015).

In each trial, the drone they operated suffered a failure and participants’ task was to decide where to crash the drone – either on a civilian site or on a military site. Crashing the drone on the civilian site led to the death of all the civilians present on the site, while crashing the drone on the military site led to the destruction of all the military material present on the site but no human death. In each trial, participants were first presented with the two possible crash sites (one at the time), followed by a decision slide presenting the two sites side by side until participants made their decision (see Figure 1). In the first part of the experiment, participants were free to decide the site of the crash (i.e., no-command condition), while in the second part they were ordered by their chain of command to protect the military material (at the expense of civilians if necessary) for strategic reasons that were not disclosed (i.e., command condition).

**Figure 1.**
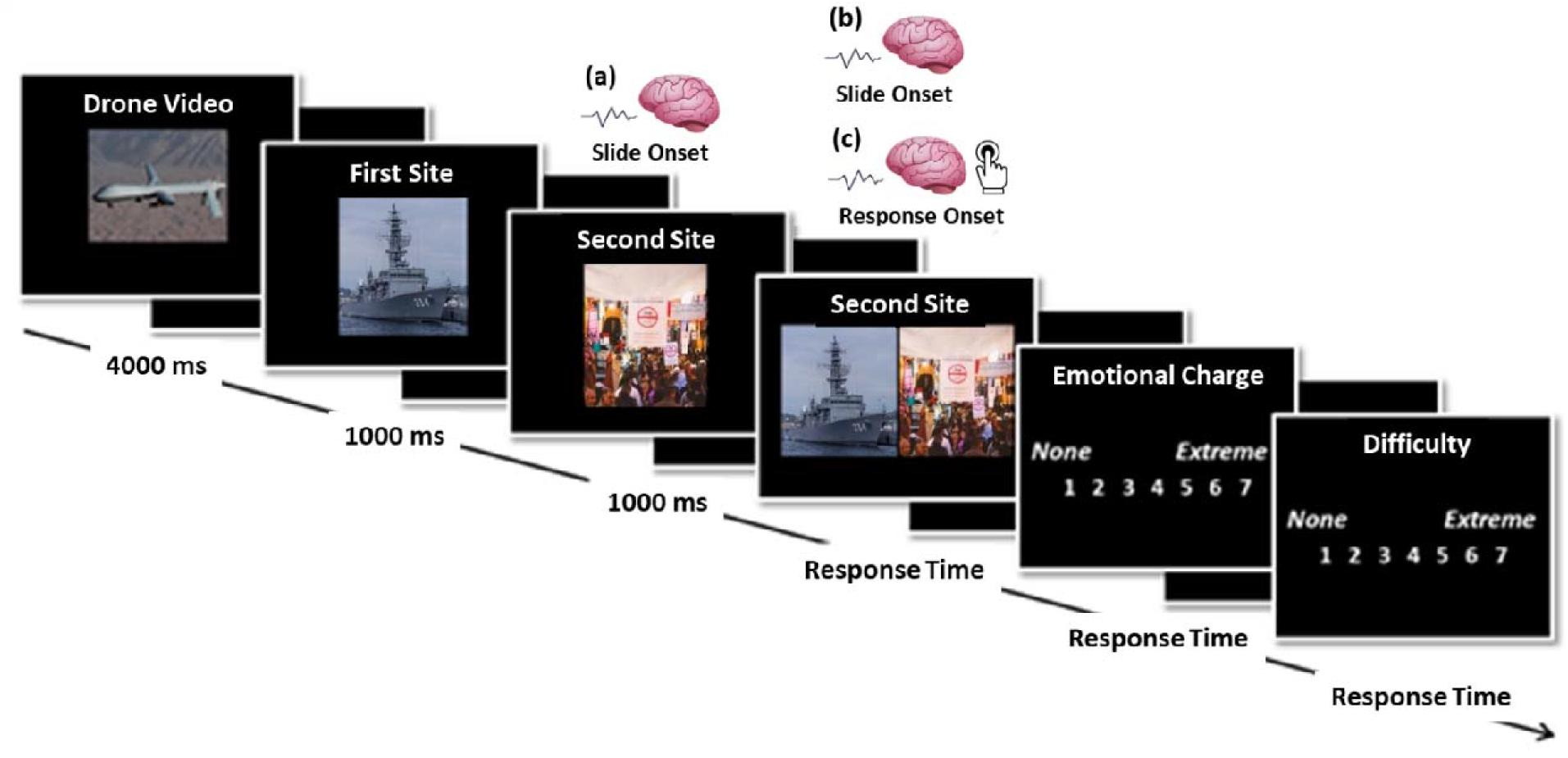
Illustration of a trial. The ERPs associated with (a) the onset of the second crash site, (b) the onset of the decision slide and (c) the response to the decision slide were investigated. The slides were separated by a 1-second white fixation point (+) displayed in the middle of a black background slide to prevent eye movements.

At a behavioral level, we investigated both the decisions and the response times observed in response to the decision slide. While the decision-making process starts right after the onset of the second option, the use of written material in previous moral decision-making studies (e.g., Sarlo et al., 2012; Pletti et al., 2015) did not allow the analysis of the ERPs associated with the onset of second option. Only the ERPs associated with the onset of the decision slide were analyzed (see section 1.2.). In the present study, the use of pictures to present the options and the relative simplicity of the experimental paradigm allowed the investigation of the ERPs time-locked to the onset of both the second option and the decision slide. While this type of analysis is seldom conducted (Gajewski, et al., 2016), we were also interested in investigating potential differences in brain electrophysiological responses as a function of the decisions made in the command mode (i.e., the moment the response key was pressed by the participants to either obey or disobey after the onset of the decision slide). In brief, at an electrophysiological level, we analyzed the ERPs time-locked to (a) the onset of the second crash site in both the no-command and the command conditions, (b) the onset of the decision slide in both the no-command and the command conditions; and (c) the response (i.e., obey versus disobey) in the command condition.

The dual-process theory provides an interesting theoretical framework to both predict and interpret the results of decision-making studies (Kahneman, 2011; Thompson, 2009). According to this theory, individuals make their decisions based on two different systems: 1) an automatic, fast, effortless, unconscious, affective, associative, and slow learning system 1 used for automatic and heuristic-based judgments and; 2) a controlled, slow, effortful, conscious and fast learning system 2 underpinning a more deliberative reasoning (Sanfey & Chang, 2008). At a behavioral level, we predicted that participants would never crash the drone on the civilian site in the no-command condition, applying the “*you shall not kill*” rule quite automatically (i.e., system 1; Chen, et al., 2009; Leuthold et al., 2015; Peng et al., 2017; Van Berkum et al., 2009). We also predicted that participants would choose to crash the UAV significantly more frequently on the civilian zone in the command condition than in the no-command-condition, as a result of hierarchical pressure.

In the case the order to protect the military material at the expense of civilian lives triggers a strong cognitive conflict between two contradicting internal rules (i.e., “*you shall not kill*” versus “*you shall obey orders*”) calling for the adoption of a more deliberative reasoning (i.e., system 2; Kahneman, 2011), we predicted that participants would take longer to make their decisions in the command condition than in the no-command condition. At an electrophysiological level, we predicted greater N200 responses to the second possible crash site in the command condition than in the no-command condition, reflecting the conflict between two opposing internal rules (i.e., “*you shall not kill*” versus “*you shall obey orders*” rule) in the command condition. We also predicted to observe greater P300 amplitudes in response to the second possible crash site in the command condition than in the no-command condition, reflecting the greater allocation of both attentional and cognitive resources to the moral decision-making process (Rigoni et al., 2010; Falco et al., 2019).

If on the contrary, receiving the order to protect the military material at the expense of human lives causes no conflict between the two internal rules and participants choose to automatically apply one of these two rules, we predicted to observe no differences in 1) response times and in 2) N200 and P300 amplitudes in response to the second site between the no-command condition and the command condition, reflecting the automatic rule-based decision-making adopted by the participants in both conditions (i.e., system 1; Kahneman, 2011). If being ordered to sacrifice innocent civilians triggers a negative affective response, we predicted greater P260 responses to decision slide in the command condition than in the no-command condition (in line with Sarlo et al., 2012; 2014; Pletti et al., 2015; Zhan et al., 2018; 2020). Finally, as the analysis of the ERPs time-locked to the response in the command condition was mostly exploratory, we made no particular predictions regarding the ERPs we would observe.

## 2. METHODS

### 2.1. Participants

33 French participants (14 females; *M_age_ =* 23 years old, *SD =* 4.45) from the University of Toulouse (France) participated in the present study. All were right-handed as assessed by the Edinburgh Handedness Inventory (Oldfield, 1971), had normal or corrected-to-normal vision and reported no history of neurological or psychiatric disorders. They received no financial compensation for their participation in the study. They were all civilian and familiar with the aeronautical and military contexts. None of them was anti-military.

### 2.2. Ethics Statement

The study was conducted in accordance with the Declaration of Helsinki (1973, revised in 1983) and was approved by the ethics committee (CERNI–Federal University of Toulouse no. 2017–040). After being informed of their rights, all participants gave their written consent.

### 2.3. Material

The present study was designed to enable the use of both the EEG and *f*NIRS techniques. The use of the *f*NIRS requires adding a 10-second baseline between the trials to ensure an appropriate signal-to-noise ratio (Pellicer & del Carmen Bravo, 2011). This long baseline between the trials significantly increases the duration of the *f*NIRS experiments. As the *f*NIRS device is likely to provoke headaches if worn for a too long period of time, we chose to present no more than 30 trials per experimental condition to the participants (as for instance in the studies of Pletti et al. 2015; Sarlo et al., 2012, 2014), which appeared as the optimal number of trials considering the specificities of both the EEG (Boudewyn et al., 2018) and the *f*NIRS (Pellicer & del Carmen Bravo, 2011) techniques.

#### 2.3.1. Norming Phase

The experimental material was composed of 30 trials of interest, in which participants had to choose to crash the drone either on civilians (i.e., civilian site) or on military material (i.e., military site), and of 12 filler trials in which participants had to choose between: 1) an inhabited site and a civilian site (four trials), an inhabited site and a military site (four trials), and 3) two inhabited sites (four trials). In order to ensure that obeying the orders in the 30 trials of interests (i.e., crashing the drone on the civilian site) would be perceived as a strong moral violation, the differences in emotional charge and severity associated with a crash had to be significantly higher for the civilian sites than the military sites.

To this aim, an on-line rating study was conducted on 150 respondents (they were recruited via social media and received no compensation for their participation in the rating study) to select the experimental material, which consisted of 30 pictures of civilian sites and 30 pictures of military sites for the 30 trials of interest; 16 pictures of inhabited sites, four pictures of civilian sites, and four pictures of military sites for the 12 filler trials. In total, 150 pictures (i.e., 60 military sites, 60 civilian sites and 30 inhabited sites) were retrieved from different free of copyright on-line databases. As it would have taken respondents too long to evaluate all 150 pictures, three on-line questionnaires, each composed of 50 pictures were created on LimeSurvey ©. For each of the 50 pictures, the respondents had to indicate on 7-point Likert scales: 1) the emotional charge (from 1 = *no emotional charge* to 7 = *huge emotional charge*) and 2) the severity of the situation (from 1 = *no severe at all* to 7 = *extremely severe*) associated with a drone crash on the site illustrated by the picture.

30 pairs of pictures associating one military site picture with one civilian site picture were created to form the trials of interest. For each pair, both the emotional charge and the severity ratings were between 3 and 3.2 points higher for the crash on the civilian site than on the military site. These differences in ratings were statistically significant with the crashes on the 30 selected civilian sites being rated as significantly higher in both emotional charge (*p* < .05) and severity (*p <* .05) than the crashes on the 30 selected military sites. For the filler trials, four civilian site pictures and four military site pictures that were not selected for the experimental trials of interest were randomly chosen and combined with 8 of the 16 pictures of inhabited sites with the lowest severity and emotional charge ratings. The 8 remaining pictures of inhabited sites were randomly combined in pairs. The fillers were added to decrease the intensity of the task, especially in the second part of the experiment, but they were not analyzed.

#### 2.3.2. Procedure

Participants were comfortably seated in the experimental room. After they signed the informed consent, a 64-electrode Biosemi© EEG cap (https://www.biosemi.com/products.htm) and a CW *f*NIRS 16-channel headband model 100 *f*NIRS system (*f*NIRS Devices LLC, Photomac MD; http://www.fnirdevices.com) were placed on their head. Meanwhile, participants had to read the written instructions of the first part of the experiment. They were explained that in this experiment they would act as a UAV operator (i.e., drone operator), gathering information on a fictive war zone in the Middle-East. In every trial, the drone they operated would suffer a failure and their task would be to decide where to crash the drone. They were told that the experiment was composed of two parts. In the first part of the experiment, participants had to decide where to crash the drone according to their preferences (i.e., no command condition). They were explained that each trial was composed of a 4000 ms drone video (see Figure 1 for the illustration of a trial), followed by a picture of the first possible crash site for 2000 ms (i.e., first site slide) and the second possible crash site for 2000 ms (i.e., second site slide). The presentation order of the type of sites was counterbalanced across trials. Finally, the two crash sites were displayed side by side until the participants made their decision (i.e., decision slide). Participants had to either press the “*a*” key of the keyboard to crash the drone on the first site – always displayed in the left part of the screen – or the “*p*” key of the keyboard to crash the drone on the second site – always displayed in the right part of the screen. After participants made their decision, they were asked to evaluate on a 7-point Likert scale: 1) the emotional charge associated with the decision (from 1 = *no emotional charge* to 7 = *extreme emotional charge*) and 2) the difficulty of the decision (from 1 = *not difficult at all* to 7 = *extremely difficult*). A 10-second inter-stimulus interval was applied to ensure an appropriate signal-to-noise ratio for the *f*NIRS signal (Pellicer & del Carmen Bravo, 2011).

Before the second part of the experiment started, participants were given the written instructions stating that for strategic reasons they had been ordered by the military hierarchy to protect the military material as far as possible (i.e., command condition). The same trials were presented both in the no-command condition and in the command condition, in a random way. At the end of the experiment, participants were debriefed and thanked for taking part in the experiment. All participants started with the no-command run. We made the unusual decision not to counterbalance the runs between the participants to prevent potential experimental biases in this specific experimental paradigm. Most people tend to be strongly influenced by the orders they receive. There was then an important risk for the participants who would have started with the command run to be still influenced by the orders (they had received in the command run) in the no-command run. It may have triggered a conflict between what they genuinely wanted to do and the orders they previously received, which may have affected the participants’ decisions, response times. This conflict may have had a much stronger influence on the ERP responses than a potential habituation effect. As presented below, stronger ERPs response were found in the command condition (i.e., second run) than in the no-command condition (i.e., first run), suggesting that the potential habituation effect (if any) may have been negligible.

### 2.4. Data acquisition

#### 2.4.1. Experimental apparatus

The experimental paradigm was presented using E-Prime 2 (Psychology Software Tools Inc., Pittsburgh, PA, USA) on a computer screen in the laboratory.

#### 2.4.2. Electroencephalography recordings

Electroencephalography (EEG) was amplified and recorded with a BioSemi ActiveTwo system (http://www.biosemi.com) from 64 Ag/AgCl active electrodes (Fp1, AF7, AF3, F1, F3, F5, F7, FT7, FC5, FC3, FC1, C1, C3, C5, T7, TP7, CP5, CP3, CP1, P1, P3, P5, P7, P9, PO7, PO3, O1, Iz, Oz, POz, Pz, CPz, Fpz, Fp2, AF8, AF4, AFz, Fz, F2, F4, F6, F8, FT8, FC6, FC4, FC2, FCz, Cz, C2, C4, C6, T8, TP8, CP6, CP4, CP2, P2, P4, P6, P8, P10, PO8, PO4, and O2) mounted on a cap and placed on the scalp according to the international 10-20 system, plus two sites below each eye to monitor eye movements. Two additional electrodes placed close to Cz – the common mode sense (CMS) active electrode and the driven right leg passive electrode – were used to drive the participants’ average potential as close as possible to the AD-box reference potential (Metting Van Rijn et al., 1991). Electrode impedance was kept below 5 kΩ for scalp electrodes and below 10 kΩ for the four eye channels. Skin–electrode contact, obtained using conductive gel, was monitored, keeping voltage offset from the CMS below 25 mV for each measurement site. All the signals were DC-amplified and digitized continuously at a sampling rate of 512 Hz, using an anti-aliasing filter (fifth-order sinc filter) with a 3-dB point at 104 Hz. No high-pass filtering was applied online. Data were analyzed with the EEGLAB toolbox (Delorme & Makeig, 2004). EEG data were re-referenced offline to the average activity of the two mastoids and bandpass filtered (0.1 - 40 Hz, 12 dB/octave), given that the low-pass filter was not effective in completely removing the 50 Hz artifact for some participants. Epochs were time-locked to 1) the onset of the second crash site in both the no-command and the command conditions, 2) the onset of the decision slide in both the no-command and the command conditions; and 3) the response (i.e., obey versus disobey) in the command condition. They were extracted for the interval between – 200 and 800 ms. A 200 ms pre-stimulus baseline was used in all analyses. Data with excessive blinks were adaptively corrected using independent component analysis. Segments including artifacts (e.g., excessive muscle activity) were eliminated offline before data averaging. A total of 6% of data were lost due to artifacts. One participant was removed from the analysis due to the poor quality of the EEG signal.

#### 2.4.3. *f*NIRS recordings

We used a procedure identical to the one used in the study of Mandrick and colleagues (2016). Due to both recording problems (i.e., the signal of eight participants was not correctly recorded) and a change in the guidelines for the analysis of *f*NIRS signal (i.e., short channels are now mandatory and the device we used was not equipped with these channels; Tachtsidis & Scholkmann, 2016), we decided to focus exclusively on the analysis of the EEG data.

## 3. RESULTS

### 3.1. No-Command Decision Mode versus Command Decision Mode

#### 3.1.1. Behavioral results

##### Decision

A Wilcoxon matched pair test was conducted on the decision rates as the data were not normally distributed (Shapiro tests: *p_s_ <* .05). Participants chose to crash the drone on the military site significantly less frequently in the command condition (*T =* 25.50, *z =* 4.46, *p <* .001; *M =* 66.84 %, *SD =* 29.94] than in the no-command condition (*M =* 95.94 %, *SD =* 8.91; see Figure 2.A.). Seven participants never followed the orders and three participants obeyed only once (~ 30% of participants strongly opposed authority). Three of them obeyed the orders more than two third of the time. 17 participants showed a proportion of obey/disobey choices between 30/70 % and 70/30 %.

**Figure 2.**
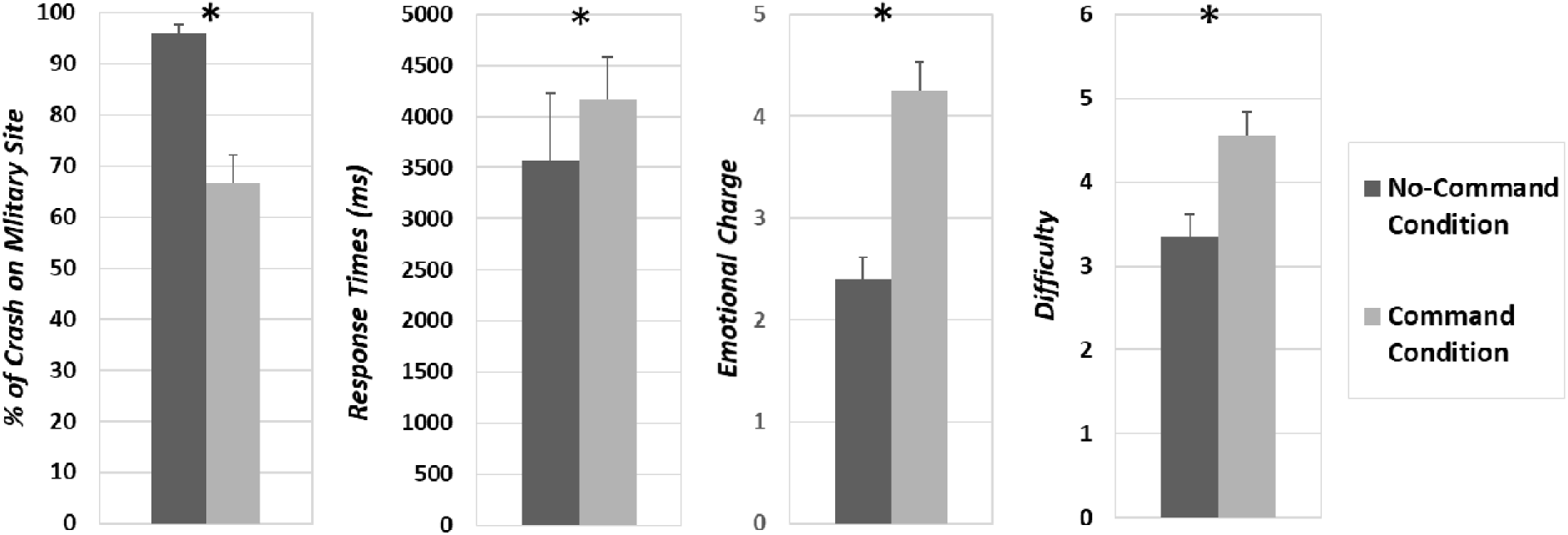
(A) Decisions, (B) response times, ratings of the (C) emotional charge and the (D) difficulty associated with the situation in the no command condition (dark grey) and the command condition (light grey). Error bars represent standard errors.

##### Response Times

A paired-samples t-test was conducted on the log-transformed response times measured on the decision slide. The analysis revealed a significant difference in response times [*t* (31) = - 3.00, *p =* .005, *CI* _95%_ (− .27, − .05); see Figure 2.B.], with faster responses in the no-command condition (*M =* 3561 ms, *SD =* 3763) than in the command condition (*M =* 4167 ms, *SD =* 2376).

##### Emotional Charge

A Wilcoxon matched pair test was conducted on the ratings of the emotional charge as the data were not normally distributed (Shapiro tests: *p_s_ <* .05). The analysis revealed a greater emotional charge in the command condition (*T =* 38.00, *z =* 4.23, *p <* .001; *M =* 4.26, *SD =* 1.55; see Figure 2.C.) than in the no-command condition (*M =* 2.40, *SD =* 1.21).

##### Difficulty of the Decision

A paired-samples t-test was conducted on the ratings of the decision difficulty [*t* (31) = − 5.52, *p <* .001, *CI*_95%_ (− 1.66, − .76); see Figure 2.D.], revealing a greater difficulty of decision in the command condition (*M =* 4.56, *SD =* 1.55) than in the no-command condition (*M =* 3.35, *SD =* 1.48).

##### Debriefing

After the end of the experiment, participants were debriefed by the experimenter. The participants who sometimes chose to crash the drone on the civilian site explained that they were afraid of the consequences of an explosion on hazardous military sites (e.g., nuclear warheads) and preferred avoiding this risk. When asked how they felt when deciding in the command condition, many participants reported that they were surprised to have felt so bad about disobeying the orders they were given, despite the fact these orders were immoral.

#### 3.1.2. Electrophysiological results: Second Site Onset (a)

Visual inspection suggested amplitude modulations between the experimental conditions for the N100, the P200, the N200, and the P300 components (see Figure 3.A.). The N100 and the P300 components were measured in terms of mean amplitude at Fz, Cz, and Pz electrodes, respectively in the 90-130 ms and the 300–430 ms time windows (e.g., Berlad, & Pratt, 1995). Due to the important amplitude differences observed before the occurrence of the P200 and N200 components, the latter were assessed in terms of peak-to-peak amplitudes at Fz, Cz and Pz electrodes (e.g., Alexopoulos et al., 2012; Falco et al. 2019). Peak-to-peak amplitudes were calculated by subtracting 1) the peak amplitudes measured in the 80–180 ms (negative peak) and the in 200–240 ms (positive peak) time windows for the P200 component (Caravaglios, et al., 2008; Falco et al., 2019; Hansch, et al. 1982); and 2) the peak amplitudes measured in the 200–240 ms (positive peak) and the 260–300 ms (negative peak) time windows for the N200 (e.g., Falco et al., 2019; Pfabigan, et al., 2011). Four 3 [Electrode (Fz, Cz, Pz)] × 2 [Decision Mode (no-command, command)] ANOVAs were performed. Post-hoc analyses were performed using HSD corrections for multiple comparisons.

**Figure 3.**
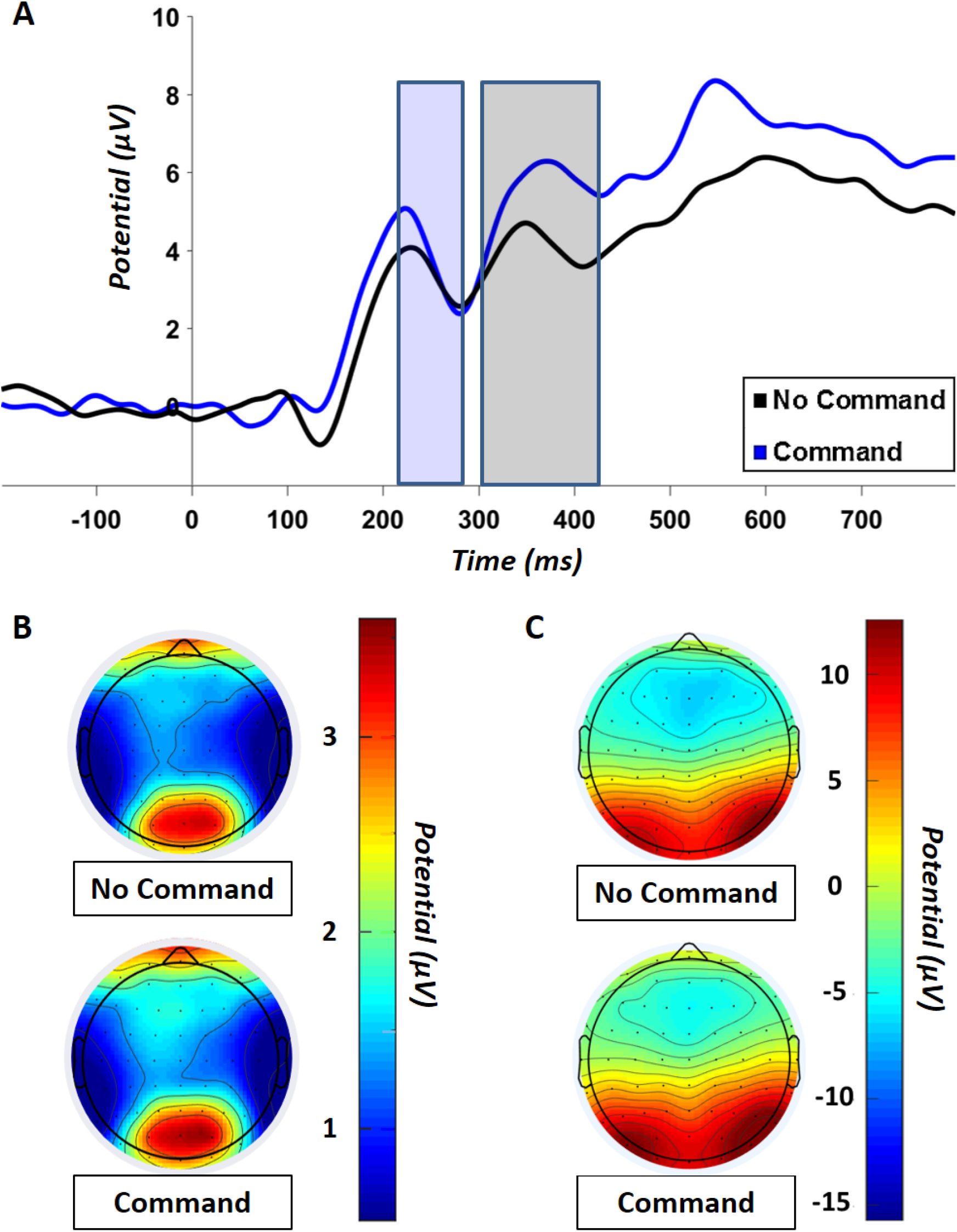
(A) Grand average ERP waveforms at Pz electrode to onset of second crash site in the no-command condition (black line) and the command condition (blue line). Scalp maps illustrating (B) the N200 peak-to-peak amplitude and (C) the P300 mean amplitude in the no-command (up) and command conditions (down).

##### N100 and P200 components

These analyses revealed no statistically significant results of interest. For the sake of clarity, they are reported in the supplementary material section.

##### N200 component

The analysis revealed a main effect of electrode [*F* (2, 62) = 6.21, *p =* .003, *ηp^2^ =* .17] with greater peak-to-peak responses measured at Fz (*M =* 2.98 μV, *SD =* 4.40, *p =* .036) and Pz (*M =* 3.25 μV, *SD =* 3.84, *p =* .004) than at Cz (*M =* 2.18 μV, *SD =* 4.03). No difference in peak-to-peak amplitude was found between Fz and Pz (*p =* .67). The analysis also revealed a main effect of decision mode [*F* (1, 31) = 38.25, *p <* .001, *ηp^2^ =* .55] with greater N200 responses found in the command mode (*M =* 4.67 μV, *SD =* 3.96) than in the no-command mode (*M =* .93 μV, *SD =* 3.33; see Figure 3.A & B.). The Electrode x Decision Mode interaction [*F* (2, 62) = .04, *p =* .96, *ηp^2^ =* .00] did not reach significance.

##### P300 component

The analysis revealed a main effect of electrode [*F* (2, 62) = 101.92, *p* < .001, *ηp^2^ =* .77] with greater P300 amplitudes measured at Pz (*M =* 4.81 μV, *SD =* 5.99) than at both Fz (*M =* − 5.88 μV, *SD =* 5.77, *p <* .001) and Cz (*M =* − 2.80 μV, *SD =* 5.98, *p <* .001); and at Cz than at Fz (*p <* .001). The analysis also revealed a main effect of decision mode [*F* (1, 31) = 6.23, *p =* .018, *ηp^2^ =* .17] with greater P300 amplitudes found in the command mode (*M =* − .45 μV, *SD =* 7.18) than in the no-command mode (*M =* − 2.13 μV, *SD =* 7.57; see Figure 3.A. & C.). The Electrode x Decision Mode interaction [*F* (2, 62) = .37, *p =* .69, *ηp^2^ =* .01] did not reach significance.

#### 3.1.3. Electrophysiological results: Decision slide onset (b)

The analyses revealed no statistically significant results. For the sake of clarity, these analyses are reported in the supplementary material.

### 3.2. Obeying versus Disobeying in the Command Condition

We also analyzed the behavioral data and the ERPs associated with the decision (i.e., obey versus disobey the order) that were made in the command condition. This analysis was conducted on the 17 participants who showed a proportion of obey/disobey choices between 30/70 % and 70/30 % in order to ensure that the number of observations of both choices would be sufficient.

#### 3.2.1. Behavioral results as a function of the decision

##### Decision

A paired-samples t-test was conducted on the data in order to ensure that the obedience and the disobedience rates were comparable. The analysis revealed that the obedience rate (*M =* 44.58 %, *SD =* 15.88) and the disobedience rate (*M =* 55.42 %, *SD =* 15.88) of the selected 17 participants were not significantly different [*t* (16) = 1.41, *p =* .18, *CI* _95%_ (− 5.48, 27.17)].

##### Response Times

A paired-samples t-test was conducted on the log-transformed response times measured when participants obeyed versus disobeyed. The analysis revealed no significant difference in response times when participants obeyed compared to when they disobeyed [*t* (16) = − .03, *p =* .98, *CI* _95%_ (− .10, .10)].

##### Emotional Charge of the Decision

A Wilcoxon matched pair test was conducted on the ratings of the emotional charge associated with the decisions to obey and disobey, as the data were not normally distributed (Shapiro tests: *p_s_ <* .05). The analysis revealed that the emotional charge was higher when participants chose to obey (*T =* 26.00, *z =* 2.39, *p =* .017; *M =* 5.21, *SD =* 2.09) than to disobey (*M =* 4.60, *SD =* 1.02).

##### Difficulty of the Decision

A paired-samples t-tests were conducted on perceived decision difficulty ratings associated with the decisions to obey and disobey. The analysis revealed no difference in difficulty [*t* (16) = − 1.89, *p =* .08, *CI* _95%_ (− 1.05, .06)] when participants obeyed compared to when they disobey.

#### 3.2.2. Electrophysiological results: Response onset on the decision slide (c)

Visual inspection suggested amplitude modulations as a function of the choice made in the command condition (i.e., obey versus disobey) for the N100 component and the P260 component. The N100 component was assessed in terms of mean amplitude in the 90–130ms time window at Fz, Cz and Pz electrodes (e.g., Berlad, & Pratt, 1995; Fabre et al., 2017). A 3 [Electrode (Fz, Cz, Pz)] × 2 [Choice (obey, disobey)] ANOVA was performed on the N100 mean amplitude. In order to investigate potential hemispherical asymmetries in the distribution of the P260 (i.e., visual inspection suggested that greater amplitudes would be found in the left hemisphere, Figure 4.B.), this component was assessed in terms of mean amplitudes in the 220-310 ms time window at F3, Fz, F4, C3, Cz, C4, P3, Pz and P4 electrodes, in order to investigate potential differences in spatial distribution (e.g., Zhang et al., 2019). A 3 [Sagittal Axis (frontal, central, parietal)] × 3 [Coronal Axis (left, medial, right) x 2 [Choice (obey, disobey)] ANOVA was performed on the P260 mean amplitude measurements. Post-hoc analyses were performed using HSD corrections for multiple comparisons.

**Figure 4.**
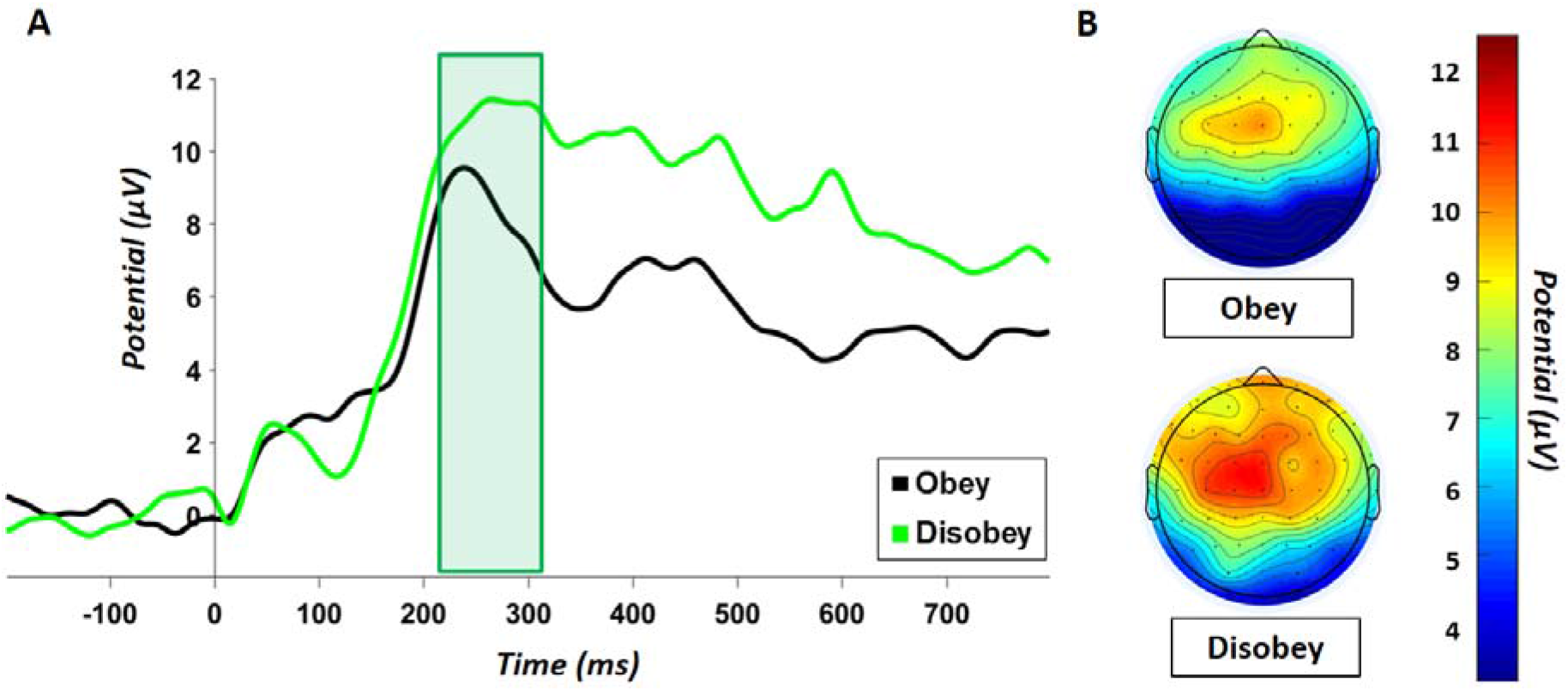
(A) Grand average ERP waveforms at Cz electrode with the choice to obey (black line) and disobey the hierarchical order (green line). (B) Scalp maps illustrating the P260 mean amplitude (220–310 ms time window) after participants chose to obey (up) and to disobey (down).

##### N100 component

This analysis revealed no statistically significant results of interest. For the sake of clarity, these analyses are reported in the supplementary material.

##### P260 component

The analysis revealed a significant main effect of sagittal axis [*F* (2, 32) = 28.61, *p <* .001, *ηp^2^ =* .64] with lower P260 responses observed at parietal (*M =* 5.17 μV, *SD* = 7.04, *p_s_ <* .001) than at both frontal (*M =* 9.09 μV, *SD =* 7.87) and central sites (*M =* 8.84 μV, *SD =* 7.79). P260 responses observed at frontal and central sites were not significantly different (*p =* .90). The analysis also revealed a significant main effect of coronal axis [*F* (2, 32) = 7.58, *p =* .002, *ηp^2^ =* .32] with lower P260 responses observed at right sites (*M =* 7.10 μV, *SD =* 7.86) than at both left (*M =* 7.89 μV, *SD =* 7.69, *p =* .018) and medial sites (*M =* 8.10 μV, *SD =* 7.76, *p =* .002). P260 responses at left sites and medial sites were not significantly different (*p =* .71). The analysis also revealed a significant Sagittal Axis x Coronal Axis interaction [*F* (4, 64) = 4.71, *p =* .002, *ηp^2^ =* .23]. Lower P260 amplitudes were found at parietal (Left: *M =* 5.58 μV, *SD =* 7.26; Medial: *M =* 5.79 μV, *SD =* 6.68; Right: *M =* 4.14 μV, *SD =* 7.16; *p*_s_ < .001) than at both frontal (Left: *M =* 8.82 μV, *SD =* 7.50; Medial: *M* = 9.38 μV, *SD =* 8.03; Right: *M =* 9.06 μV, *SD =* 8.30) and central sites (Left: *M =* 9.27 μV, *SD =* 7.99; Medial: *M =* 9.14 μV, *SD =* 8.11; Right: *M =* 8.11 μV, *SD =* 7.42) along the whole coronal axis. Lower P260 amplitudes were also found at right sites than at both medial and left sites, both at central (medium: *p =* .037; left: *p =* .012) and parietal sites (medium: *p* < .001; left: *p <* .001). Finally, the analysis revealed a significant Sagittal Axis x Choice [*F* (2, 32) = 5.39, *p =* .010, *ηp^2^ =* .25; see Figure 4.A & B] with greater P260 responses observed at frontal, central and parietal sites when participants chose to disobey (Frontal: *M =* 9.82 μV, *SD =* 9.62; Central: *M =* 10.20 μV, *SD =* 9.09; Parietal: *M =* 6.89 μV, *SD =* 7.66) compared to when they chose to obey (Frontal: *M =* 8.36 μV, *SD =* 5.61, *p =* .023; Central: *M =* 7.49 μV, *SD =* 6.00, *p <* .001; Parietal: *M =* 3.45 μV, *SD =* 5.94, *p <* .001). Lower P260 amplitudes were also found at parietal sites (*ps <* .001) than at both frontal and central sites for both types of choice. The main effect of choice [*F* (1, 16) = 1.84, *p =* .19, *ηp^2^ =* .10] and the Coronal Axis x Choice [*F* (2, 32) = .63, *p =* .54, *ηp^2^ =* .04] and the Sagittal Axis x Coronal Axis x Choice [*F* (4, 64) = .57, *p =* .69, *ηp^2^ =* .03] interactions did not reach significance.

## 4. DISCUSSION

The present experiment aimed at investigating the impact of hierarchical pressure on subordinates’ decision-making and the associated electrophysiological neural correlates. Participants acted as UAV operators and had to decide to crash their defective drone either on a civilian site killing all civilians present on the site, or on a military site destroying the site’s military material but preventing human losses. They had to make their choice either by themselves (i.e., no command condition) or after receiving the order to crash the drones on civilian sites (i.e., command condition).

In the no-command condition, participants almost always chose to crash the drone on the military equipment and save the civilians (i.e., about 96 % of the time). During the post-experimental debriefing, the participants who sometimes chose to crash the drone on civilian sites explained that they were afraid of the consequences of an explosion on hazardous military sites (e.g., nuclear warheads) and preferred avoiding this risk. In the command condition, participants chose to obey orders and sacrifice civilians to protect the military material one third of the time (in line with the variation of Milgram’s experiment in which the experimenter has not physically present in the room; see Perry, 2013).

In the command condition, participants took longer to make their decisions and reported greater perceived difficulty and emotional charge associated with the decision than in the no-command condition. Greater N200 amplitudes were also found in response to the second site picture in the command condition than in the no-command condition (a). This result suggests that deciding in the command condition was more conflictual for participants than deciding in the no-command condition, supposedly reflecting the conflict between two competing internal rules: “you shall not kill” versus “you shall obey orders” (Gehring & Fencsik, 2001). Greater P300 responses were also found in response to the second site picture in the command condition than in the no-command condition. The decision process appears to have required greater attentional and cognitive resources in the command condition than in the no-command condition (Gray et al., 2004; Linden, 2005). Taken together, the present results suggest that overall participants mobilized greater attentional and cognitive resources to resolve the conflict between two competing internal rules when they had to decide under coercion compared to when they had to decide by themselves (i.e., no command condition) and automatically applied the “you shall not kill” rule (Kahneman, 2011; Sanfey & Chang, 2008). Moreover, no difference in response times was found as a function of the decision in the command mode, suggesting that the decision to obey was not more automatic than the decision to disobey. To sum up, receiving immoral orders appears to have increased the cognitive effort associated with the decision-making and did not result in an automatic rule-based decision-making.

A complementary analysis (c) was conducted on the ERPs time-locked to the decision made in the command condition on the data of the participants who showed a proportion of obey/disobey decision ranging from 30/70 to 70/30 percent. Greater P260 amplitudes component (i.e., late P2-component) were found after participants chose to disobey than to obey orders. In various studies where participants had to decide on moral dilemma situations, greater P260 responses were interpreted as reflecting an aversive affective reaction and was found to be positively correlated to the to the unpleasantness experienced during moral decision-making (Sarlo et al., 2012; 2014; Pletti et al., 2015; Zhan et al., 2018; 2020). Taken together these results suggest that the greater P260 amplitudes observed after participants disobeyed may reflect a greater negative affective reaction to disobedience than to obedience, even though obeying implies sacrificing innocent civilians.

As individuals, when presented with moral dilemmas, tend to choose the option that minimizes the intensity of negative emotions associated with the decision (e.g., Pletti et al., 2016), the results of the present study show that subordinates may sometimes choose to obey immoral orders, not only because coercion affects their sense of agency (Milgram, 1974; Caspar et al., 2016), but also in order to avoid the negative affective response associated with disobedience. Many psychologists, including Milgram himself (Milgram, 1963; 1965), were caught by the extreme emotional distress of the participants who took part to the *obedience to authority* experiments when they chose to obey the experimenter’s orders (e.g., Baumrind, 1964; Miller et al., 1995; Perry, 2013). This observation raised the following question: why would some individuals prefer going through such an affective distress instead of “simply” disobeying? Yet, the possibility that disobeying malevolent authority could trigger an even greater emotional distress than obeying immoral orders, at least in some individuals, has never been considered.

This greater affective response to disobedience can be explained in two ways. On the one hand, as disobeying orders is usually harshly punished by the authority in charge (Bocchiaro & Zamperini, 2012), especially in the military (Gaeta, 1999; Munro, 2019), individuals may anticipate the fear and the stress of being punished (Gray, 1987). This hypothesis is in line with the results of previous studies, where greater P2 responses were found when participants fearer and anticipated a painful punishment (e.g., Lei et al., 2019). However, we assume that it is unlikely that participants feared being punished in the present experiment, because disobedience was not sanctioned. On the other hand, Graham and colleagues showed that people (especially conservative ones) consider disobedience to authority a moral violation (Graham, et al., 2009, 2011).

. Moral violations trigger negative self-conscious emotions (Tangney et al., 2007), such as shame, guilt, or embarrassment (Fabricius, 2004; Lee, 1999), and were found to be associated with greater P2 brain responses (e.g., Chen, et al., 2009; Gan et al., 2016; Leuthold et al., 2015; Peng et al., 2017; Van Berkum et al., 2009; Zhu et al., 2019). This negative affective reaction observed in response to moral violations is thought to play an important role in moral decision-making, as it may serve the adaptive purpose of promoting people’s compliance to moral and social rules (Blair & Fowler, 2008; Tangney et al., 2007). Taken together, the results of these previous studies suggest that the greater P260 responses observed after participants chose to disobey (to save the civilians) may reflect the fact that disobeying immoral orders constitute a greater moral violation than obeying them and sacrificing innocent civilians.

While a large part of the literature has focused on the dire consequences of obedience to authority (Milgram, 1963; 1965; 1974), some authors have highlighted its positive side arguing that obedience is essential to the effective functioning of human society (Darley, 1995; Ent & Baumeister, 2014). Humans have evolved to participate in complex cultural systems, whose success highly depends on the capacity of the individuals composing this system to coordinate their effort to pursue a common goal (Baumeister, 2005). This need for coordination calls for a hierarchical organization of the group, with authorities providing both leadership and protection; and subordinates obeying and respecting authority. Well-organized groups (characterized by great leadership and a high level of obedience) are able to grow in size and endure, while less organized groups (characterized by a poor leadership and a strong tendency to disobey) tend to be absorbed or destroyed by the formers (Fukuyama, 2011). As obedience plays an important role in maintaining the stability of a group, individuals “equipped with an affective system” preventing them for disobeying authority may have been favored by natural selection, explaining why disobeying authority is so difficult for many people.

Interestingly, in both the present study and the study of Milgram (1963), some participants chose to strongly oppose authority (respectively, 30% and 35% of the participants). Extremely rigid groups with extreme levels of authority and obedience very hardly adapt to changing conditions and are likely to quickly dissolve, especially when authorities perform poorly and are more interested in maintaining/expending their own power than in guaranteeing the group’s well-being and stability, thus jeopardizing its chances of survival (Acemoglu & Robinson, 2012; Ent & Baumeister, 2014). While a group formed in large part by obedient individuals tend to be well-organized and have a greater chance of survival, the presence of some individuals who have the capacity to oppose authorities (supposedly due to a decreased affective reaction to disobedience) when the latter work against the general interest may also constitute an important safety net for the preservation of the group (Boehm, 2012). These “disobedient” individuals may be fewer in number (compared to the obedient ones), on the one hand because opposing authority comes with a very high cost (i.e., resource limitation, ostracism or even death; Bocchiaro & Zamperini, 2012), which lowers their chances of survival and have offspring, and on the other hand, because too many disobedient people in a group may jeopardize its stability.

Further investigation is now necessary to confirm that disobedience to authority triggers a negative affective response in (some) subordinates, who might choose to obey in order to avoid this aversive reaction. Conducting this same experiment using the *f*MRI technique would enable to further investigate the neural bases of both obedience and disobedience to authority and help confirm the important role played by affective processes in moral decision-making under coercion. We proposed two possible explanations for the affective distress experienced by participants after they disobeyed: 1) the fear triggered by the anticipation of retaliations and 2) the negative self-conscious emotions triggered by the violation of the obedience moral rule. Further research is needed to either rule out one of these explanations or determine to what extent each of them accounts for the affective distress experienced by participants after they disobeyed. It is highly probable that important inter-individual differences will be found; with 1) egoistic traits predicting for a greater fear of retaliation; and 2) agreeable dispositions (which are positively correlated to the tendency to both conform to social groups and avoid upsetting others; DeYoung et al., 2002; Roccas et al., 2002) predicting for a greater susceptibility to negative self-conscious emotions (possibly explaining why agreeable people are prone to obey malevolent authority; Bègue et al., 2015). We were not able to demonstrate that the participants who always disobeyed were less sensitive to the negative affective reaction to disobedience. It would be interesting to investigate this question and to test whether this decreased sensitivity is due to 1) a lower affective distress reaction compared to more obedient individuals, and/or that 2) a greater capacity to regulate these negative emotions (Goldin et al., 2008; Grecucci et al., 2015). Finally, if people obey malevolent authority to avoid the affective distress associated with disobedience, it may be interesting to investigate whether training them to apply emotion regulation strategies – such as emotion suppression or emotion reappraisal; Goldin et al., 2008; Grecucci et al., 2015) – could increase resistance to malevolent authority.

